# An analysis of the full song of five free-living urban European Blackbirds *(Turdus merula*); a network approach

**DOI:** 10.1101/505206

**Authors:** Koenraad Kortmulder

## Abstract

Is the song repertoire of male blackbirds safe-guarded against loss of variability during the breeding season? In order to answer this question the repertoires of five males were analysed from the viewpoint of network theory. In four of the five males a strong coherence of the repertoire was found to exist in anastomoses between strophe types, same elements being reached from different beginnings. The succession of strophe types in song sessions betrayed a mixture of relatively predictable cycles and chaotic connections. The former should facilitate repeated retrieval of a considerable part of the repertoire, while the chaotic component keeps the whole repertoire readily accessible from any point in a singing session. The same mixture may be considered optimal for binding the attention of conspecific listeners. It is concluded that the existing structure of blackbird song repertoires is favourable for the conservation of its richness, but it is as yet uncertain whether this is due to evolutionary (*i.c.* sexual) adaptation.

## Introduction

The song of Turdid birds has been a favoured object of study through the past half century, partly because species range from relatively simple to extremely complex song structures. Whitney (1981a,b) chose the Varied Thrush^1^ because of its relatively small repertoire and small number of song parameters. The average number of its song types is five and pitch and modulation frequency suffice as parameters distinguishing song types. Among the richest in song performance are the American Robin, the European Song Thrush, the European Blackbird and the Nightingale^2^. Dobson & Lemon (1979) compared the songs of five North-American thrushes which differ greatly in song variability. The present paper is about European Blackbirds.

Blackbirds are widely known among European town inhabitants because of their bold behaviour and beautiful voice. Many studies of their song have been made by *e.g.* Messmer & Messmer (1956), Thielke-Polz & Thielke (1960), Hall-Craggs (1962), Todt *(e.g*. 1970, 1975), Todt & Wolffgramm (1975), Ripmeester (2009). All authors agree that a typical blackbird strophe consists of a *melody* or motif part followed by a higher-pitched *shrill*. No more than a hint is given by some that a second melody, and sometimes a third, may follow the first shrill. Second melodies were common in my birds (range 34-63% of the strophes) in *both* ‘tradition groups’ that could be distinguished (below). Successive strophes are separated by silent intervals of at least a few seconds. Series containing up to hundreds of strophes alternate with lengthy flying-away intervals in which the bird feeds or interacts with enemies, rivals, mates or young. The longer series tend to occur at dawn and in the evening. Early in the season, singing is practically confined to these times. It is gradually extended to the whole day later.

The complexity of the song in some turdid species may be taken as an indication of sexual selection, comparable to the peacock’s tail or other extreme visual displays of the male sex. If so, good singers should be found to sire more young than poor ones. Should song be the only way in which males distinguish themselves, such reproductive advantage would be expected especially in second broods, because of the limited time devoted to singing early in the season. If all this is true, it is important for a male to acquire and maintain a rich repertoire of song types and permutations of the same. The principal question of this paper is therefore: *how do males manage to not let their performances impoverish with time*. Just imagine: if you or I had a repertoire of, say, 30 tunes that we hum spontaneously from time to time, we would be likely to lose at least some of them within a month or two unless, for instance, we had them written down or always hummed them in the same fixed order, preferably with a cue of simple scale relationships between them such as each tune a fifth lower or a fourth higher than its predecessor.

Diverse methods have been used for the statistical analysis of complex song. Todt (1970, 1975) and Whitney (1981b) developed *cybernetic models*, with the appropriate negative feedbacks counteracting immediate repetition of a song type or its being followed by similar types. Todt’s model was tested in the computer by Todt & Wolffgramm (1975) and found to produce results comparable to normal blackbird song. Whitney (1981a,b) did the same for the Varied Thrush but admits that other models are possible. Todt & Hultsch (1998b) found *hierachical patterns* in the memorisation and retrieval of song types in the Common Nightingale. A theoretical study of hierarchical configurations in behaviour by Dawkins (1976) was criticised by Nelson (1989), who argued that both the theoretical and the real world are too complicated to be described by hierarchical or Markovian models alone. As an example from reality he analyses the song of the European Song Thrush in great detail and shows how Markovian chains and hierarchies are closely intertwined.

*Markov chain analysis* has been advocated by Dobson & Lemon (1979) and others (overview in ten Cate & Okanoya, 2012). A very special way of how birds sequence song types has been reported by Nelson (1973) in Swainson’s Thrush^3^. Successive songs of this bird seem to be related to each other by harmonic relationships, song *n* being followed by *n* + 1 2 steps up or 3 steps down in a roughly pentatonic scale. His findings have been criticised by Dobson & Lemon (1979) who could not identify key-notes in the songs of several individuals of the same species and, when using an alternative method, found similar but not precisely the same patterns of succession Nelson found. However, as Whitney (1981b) remarked, this is insufficient reason to reject Nelson’s finding^4^.

Studies on blackbirds comprising a male’s whole repertoire are few. Hall-Craggs (1962) followed a male through an entire singing season. She documented how strophes developed with time from single motives of a few notes. Morgan *et al.* (1976) applied cluster analysis to her data and disclosed several preferred sequences of strophe types within series. Hesler *et al.* (2012a,b) measured repertoire size as the number of different note types. The blackbirds I studied exhibited a very few instances where one or two notes seemed to be thrown in between melodies (see also variations, below), but clearly the melody was the main smallest unit of recombination. So I focussed on whole melodies. Detailed analyses of song structure comparable to those by Isaac & Marler (1963) on the Mistle Thrush^5^ and others mentioned above have not been done with blackbird song. The present paper may supplement this.

In this paper I adopt a network approach, since this seems the most appropriate to the main question of how a male blackbird keeps the richness of its repertoire up.

## Material and Methods

Four fully adult male blackbirds and a presumably first-spring male were studied in and around the gardens of Rijn- and Schiekade 111-116, Leiden, The Netherlands. Recordings were made during evening song when the season was well advanced, most of them in June or July. Three males, recorded in 1974, 1976 and 1978 respectively shared some melodies, whereas the fourth adult male and the youngster, both recorded in 1985, shared some others between them and clearly belonged to one, different, song tradition. By that time none of the shared melodies of the other three were heard again.

For analysis, I selected series with least disturbance and containing at least 125 strophes in at most two uninterrupted runs. Exceptions had to be made for the 1985 adult male whose performance came in bits and pieces with a longest pause exceeding 90 seconds summing up to 145 strophes, and the yearling who yielded only 70 strophes in all.

I used a 4000report-L UHER tape recorder with the microphone fixed at the focus of a polyester paraboloid mirror. Lacking the supreme musical talent of Joan Hall-Craggs, who recorded blackbird song directly on staved paper, I depended on two methods to translate acoustic into visual patterns: (a) feeding the signal into a Siemens Oscillomink S ink-jet recorder via a digitalising interface or (b) the computer programme ‘Audacity’ which translated the sound in spectrographic screen images which were subsequently printed. The sonograms produced with the former method had the advantage that they could be easily superimposed with transparent paper, showing amazingly perfect mapping of same type songs in both pitch and rhythm (Messmer & Messmer, 1956, *pp.* 377-9). The latter procedure proved just a little more perfect in the identification of some heavily disturbed songs (background noise of trains or planes passing). All series were finally checked with method (b).

All strophes that began with the same opening melody — they never start with a shrill — were assigned to the same strophe type, as other authors have done. Strophe types were indicated by a figure code. Often, opening melodies came in two or more variations, which I classified as a, b, c, etc. For such variations to be assigned to the same melody type, they had to have at least the first two or three notes in common. In some cases I ignored the presence or absence of a small opening hiccup. Most variations consisted of omitting, inserting or doubling one note. Repertoire size was defined as the number of different strophe types, lumping variations. Add those melodies which never stood at the beginning of a strophe and it will be clear that the array of melodies of a bird can be much larger than its strophe type numbers (table 1).

**Table 1.** Some parameters of the song repertoires of five blackbirds. Convergent = arrows converging on melody (red in figs. 3); divergent = arrows diverging from melody and either ending on next melody (upper rows) or on any next element (lower rows), black in figs. 3; complexes = groups of strophe types bound together by connections of succession; continents = very large complexes; islands small complexes; mels = melodies; second mels = percentage of strophe types with more than one melody. The numbers of different opening melodies would be considerably higher if melody variations (a, b, c etc.) were counted separately.

|  | ♂yng | ♂Sleg | ♂tuin | ♂Sand | ♂1974 |
| --- | --- | --- | --- | --- | --- |
| **Strophe type anastomoses** |  |  |  |  |  |
| convergent (red) | 21 | 31 | 26 | 22 | 11 |
| total | 36× | 61× | 47× | 37× | 24× |
| idem to potential opening melody | 11 | 14 | 19 | 11 | 4 |
| total | 15× | 36× | 36× | 19× | 12× |
| divergent (black) melody to melody | 5 | 6 | 4 | 18 | 11 |
| total | 12× | 24× | 19× | 82× | 56× |
| divergent (black) melody to any | 13 | 16 | 14 | 28 | 27 |
| total | 50× | 83× | 86× | 149× | 129× |
| **Complexes** |  |  |  |  |  |
| ‘continents’ | 1 | 1 | 1 | 3 | 0** |
| ‘islands’ | 0 | 3 | 7 | 5 | 22 |
| summed | 1 | 4 | 8 | 8 | 22 |
| **Melodies** |  |  |  |  |  |
| different opening melodies | 7 | 29 | 28 | 33 | 33 |
| other melodies | 3 | 22 | 13 | 30 | 26 |
| summed | 10 | 51 | 41 | 63 | 60 |
| second melodies*** | 63% | 52% | 54% | 52% | 34% |
| sample | 70 | 134 | 145 | 209 | 160 |

\* might be an underestimate if shrills of male yng were identifiably diverse, which is uncertain.
\*\* or one à two very small ones.
\*\*\* with or without a shrill in between (almost never in male yng).

Intervals between strophes varied considerably in length, but there was practically never room for doubt where one strophe ended and the next began. For this study I focussed on the melody parts of the strophes, leaving aside interval lengths and details of shrills. Only, it may be said that the shrill or shrills typically ‘belonged’ to the preceding melody of the same strophe, even where several diverse shrills could follow the same melody. Different melody variations were *usually* followed by different shrills. Particular attention was given to internal structures at the melody and strophe levels (section b) and to the sequence of strophe types in a singing session (section c).

In the terminology of network theory, units — whether strophe types or parts thereof — are called *nodes*. If connections between nodes are one-way, as in the present case, the former are named *arcs*; the whole network representation a *digraph* (directional graph). Nodes connected by more than two arcs are *hubs*. The *degree* of a hub is the number of connecting arcs beginning or ending on it (Anonymous, n.d.; Douw, 2010; de Haan, 2014).

An important parameter in the analysis of networks is L, the shortest path length (in number of arcs) between two nodes, averaged over all pairs of nodes. In practice, one gets a good impression of the value of L by assessing the distance of 5 or 10 randomly chosen pairs. An alternative method is to count the number of steps necessary to reach, from a randomly chosen node, all others. For practical purposes, I set the limit at 90%. Both methods were applied for all four full-grown males. As a rule of the thumb, when L is smaller than the ^*e*^ log (ln) of the number of nodes, the network may be qualified as a “small-world” network. This means that any node can be reached from any other in a very small number of steps (Watts & Strogatz, 1998)^6^.

## Results

### a. Repertoire contents

Let me first introduce the five males and their repertoires (figs. 1A-E). They are named: ♂1974, ♂Sand, ♂Sleg, ♂turn and yng. As already mentioned, I have categorised strophe types according to the first melody of the strophe. Variations in the opening melody or further on were ignored. Extensive attempts to include second and third melodies in the analysis made results less rather than more clear; so I stuck to the former criterion. On the abscissa are the *strophe types*, arranged in descending order of frequency of occurrence. Evidently, the birds differ as to the total number of strophe types (*N_t_*): from 7 in ♂yng to 33 in ♂Sand and ♂ 1974. On the ordinate are the *frequencies* with which they occurred. *N_S_* = number of strophes in the sample.

**Figure 1.**
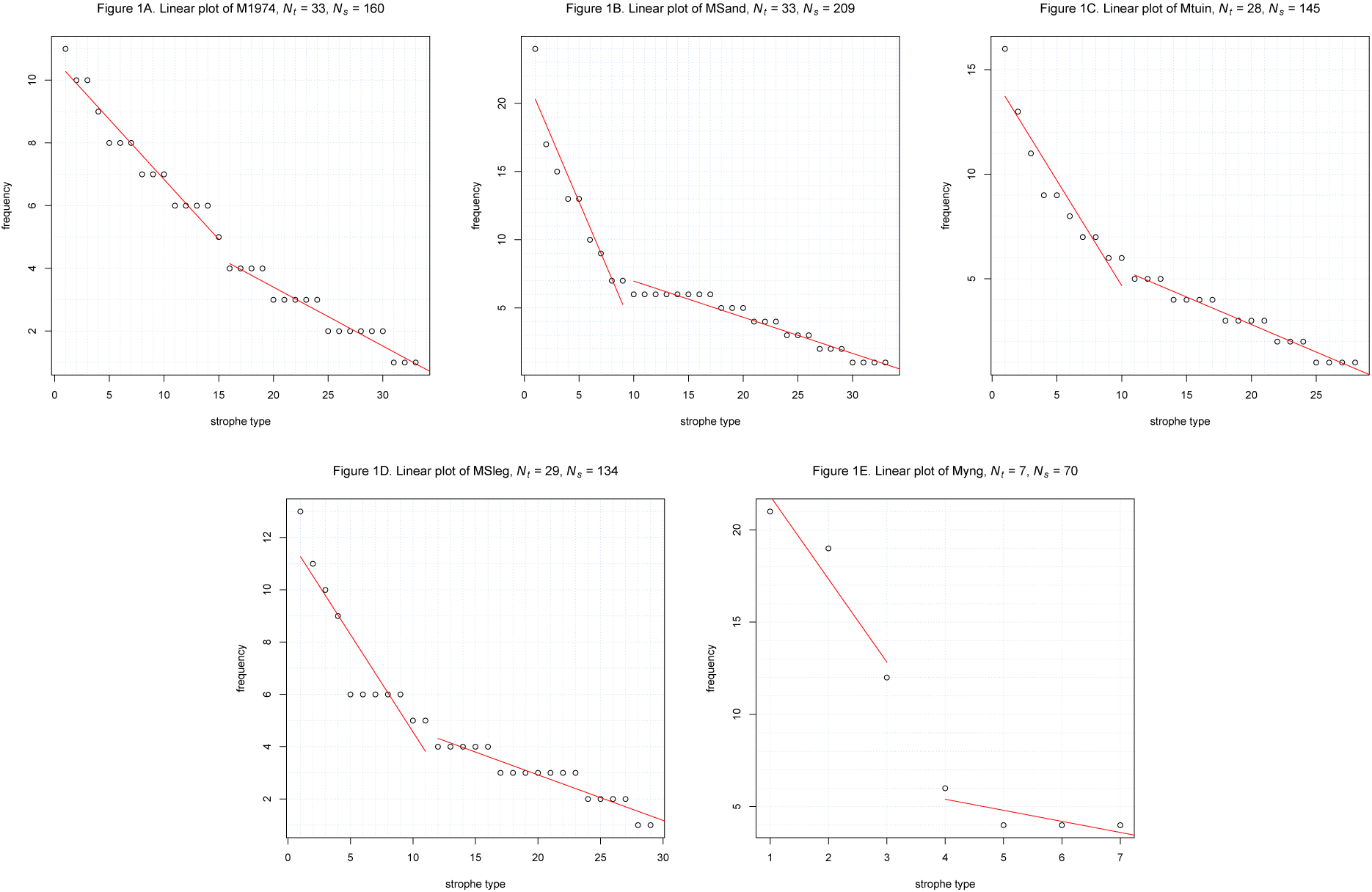
Frequencies of occurrence of all strophe types in the repertoires of males 1974, Sand, tuin, Sleg and yng. Types in order of decreasing frequencies. Red lines indicate the most probable position of the kink.

As already noted by Todt (1967a,b) strophe types differ in frequency of occurrence. In addition, at least in each of the four adult birds, there is a kink in the curve suggesting two populations of strophe types, those of the most frequent types being disproportionally well represented (preferred motifs according to Dabelsteen, 1984). The most likely position of the kink has been calculated with the ‘Multiple changepoint test using binary segmentation’ (red lines in the figures).

Figs 2A-E show the same data with a logarithmic ordinate. The reason for this transformation is that Nelson (1989) found a so-called Zipf relationship for the frequencies of song types of the European Song Thrush, that is: the double log graph yields an approximately straight line, a sort of relationship also existing in human language. Apparently, ‘Zipf’ does not apply to the blackbird song, at least not at the strophe level. Instead, the left part of the graph (most frequent strophe types) tends to be exponential (straight in the log transform), the right part straight. The latter, by the way, suggests a rather closed collection of strophe types, or in other words there is no tail of rare types. at least during each of the singing sessions under study. Other series of the same birds would have to be analysed to be sure that there are no ‘hidden’ strophe types. Dabelsteen (1984, *p.* 235) states that dawn and evening song contain the same notes and melodies, but gives no details.

**Figure 2.**
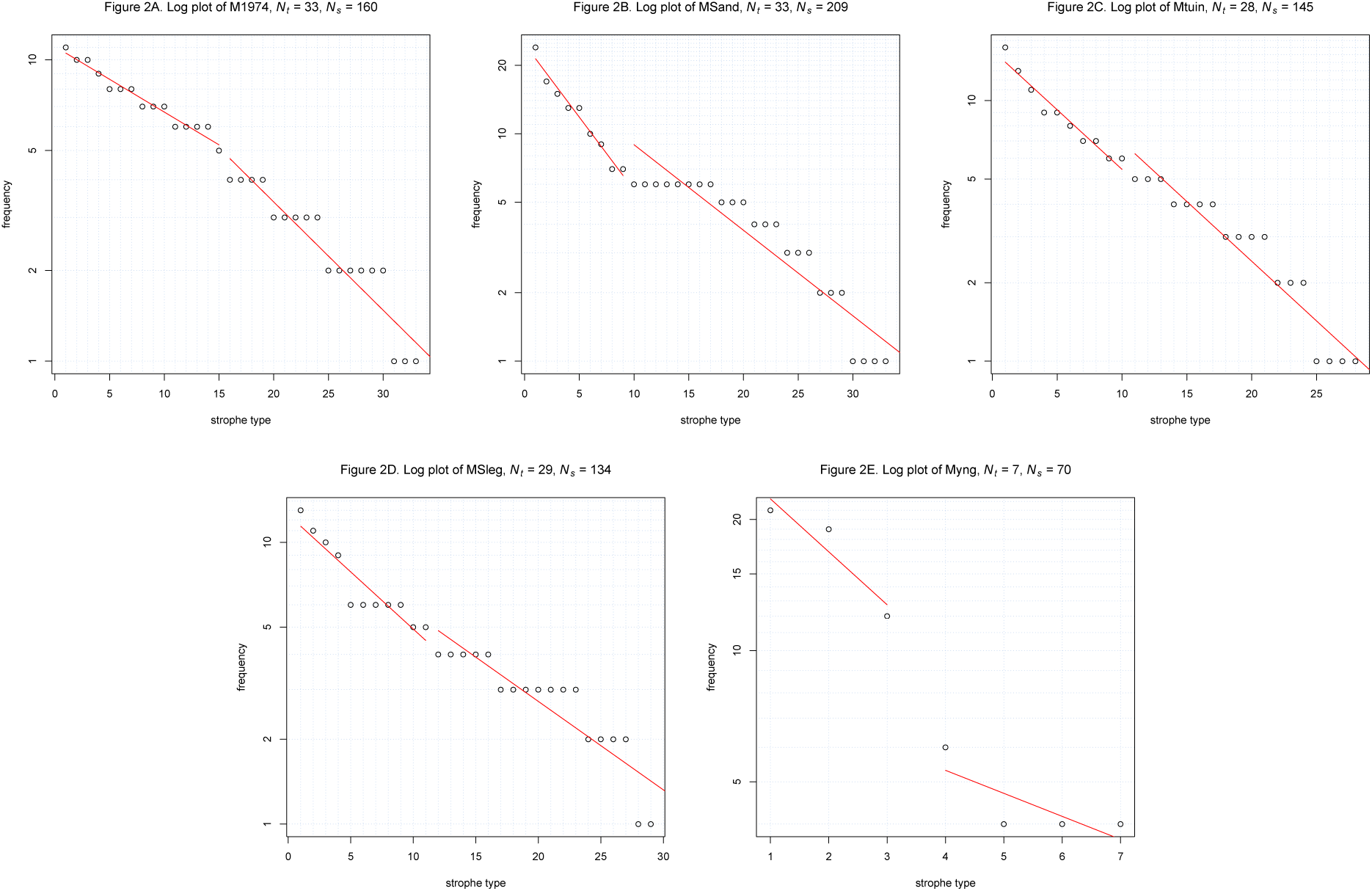
As figs.1, ordinate in log transform.

At this point there is no obvious reason for the dual nature of the repertoire. I shall come back to it in section c on strophe type sequencing.

### b. Strophe stucture and interstrophe connections

Take the reper-toire of ♂1974 as an example. (Figures 3A-E are on *p.* 17). Figure 3A represents all of its melodies and shrills and their connections by succession within strophes. Melodies and shrills are linked by predominantly divergent connections, that is: identical melodies may be followed by different shrills followed by different melodies *etc.* If one expects that repertoire stability depends on the density of connections *between* strophe types — which is a corollary of network theory: the more connections, the smaller the risk that the network will become fragmented — figure 3A must come as a disappointment. 16 of the 33 strophe types appear as islands, unconnected with other types; eight are connected pairwise. Most conspicuous exceptions are the small webs surrounding the 33 and the 222^2^ melodies, both of which are to the human ear very prominent members of the repertoire. (A third prominent member, 51, doesn’t distinguish itself here). Note that this is the male with the largest number of strophe types and melodies of all (Table 1)! Shouldn’t he be expected to have the most stable system, with the largest density of connections? Or are islands the rule?

**Figure 3.**
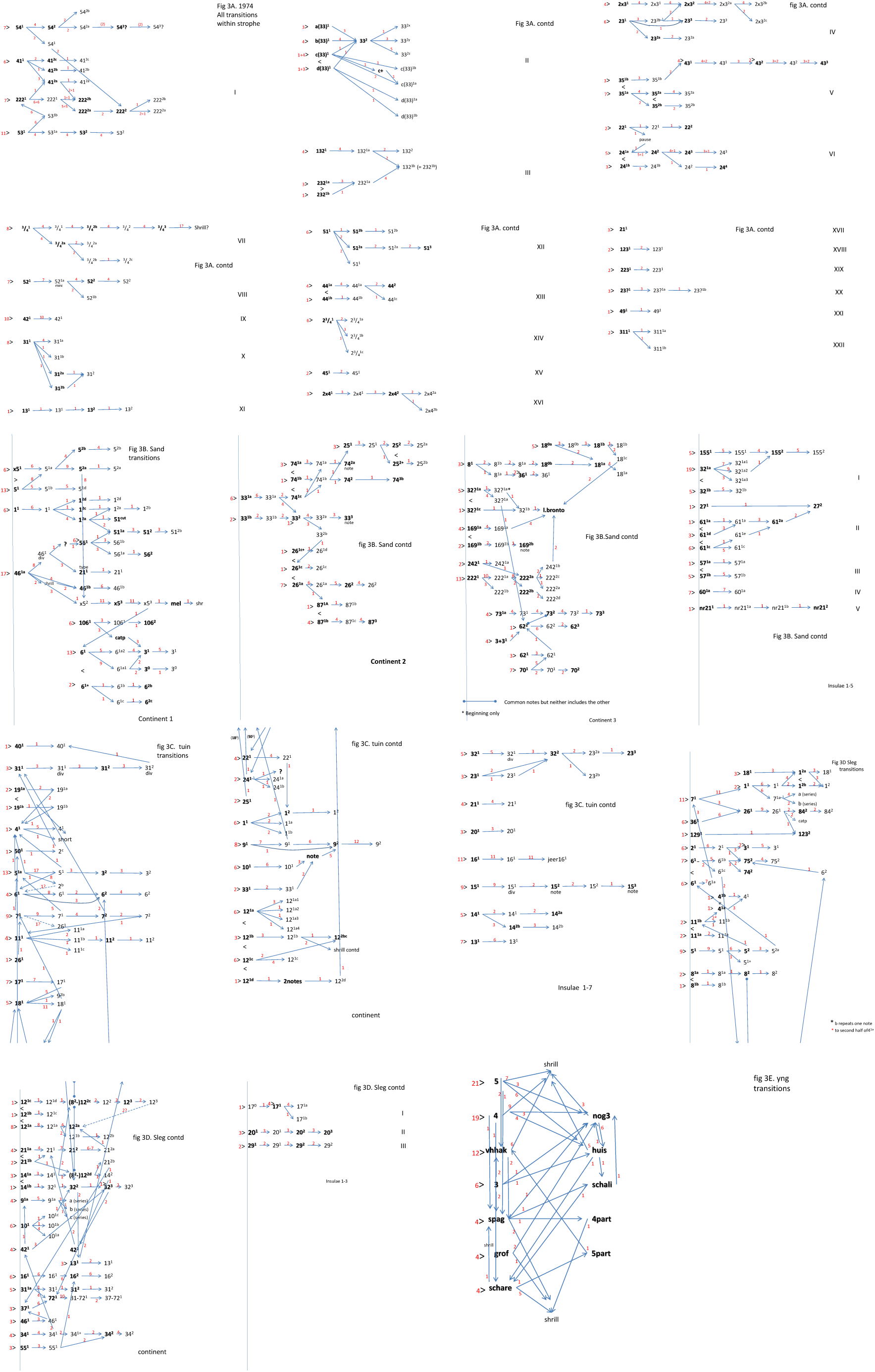
A-E. All transitions between elements within strophes in males 1974, Sand, turn, Sleg and yng. Melodies are in bold print to distinguish them from shrills. Frequencies of transitions and starts in small, red figures. Roman figures indicate islands.

As to the latter question: no, the graphs of the other four males abound with interconnections between strophe types, so much so that the great majority of strophe types are bound together in one or a very few ‘continents’ and only a small number of ‘islands’ (figs 3B-E and table 1). Champion is the young male: (fig.3E). His four melodies that only occur at the beginning of strophes (5, 4, 3 and grof) may be followed by respectively four, four, four and one different second melodies^7^. Three that are sung only later in the strophe (nog3, huis and schali) are preceded by seven, four and one different ones^8^.

This is curious; or could the density of interconnections between strophe types be a property of young, beginning singers, and would it gradually diminish as the bird becomes more accomplished? Look at table 1 again. The five males can be read as representing a series from very young to very mature singers, beginning with the youngster and ending with ♂ 1974. The actual ages of the males are unknown, the identification of one of them as a beginner being the most sophisticated guess, but all parameters in the table taken together do support a directional relationship as follows: the number of strophe types roughly increases from left to right and so does the total number of melodies in the repertoire; the strophe types lose more and more connections between them, resulting in an ever larger number of ‘islands’. Concurrently, the percentage of strophes with second melodies decreases.

#### di- and convergences

The connections between strophe types may result in convergences (same melodies being preceded by any of two or more others). The relevant arrows in figures 3 are marked in red. As may be seen in table 1, the number of red arrows per network decreases from left to right, with the exception of male yng (who has less melodies in total). The same trend may be seen in the numbers of those red arrows that connect to types of melodies which can also be found at the beginning of strophes, with the exception of ♂tuin, where this number peaks at 19. ♂tuin’s high value for this kind of red arrows corresponds to his low score for the number of melodies which exist only as second in the strophe (‘other mels’ in the table) and to his relatively low score in divergences (table). On the other hand, diverging patterns increase in the table from left to right. ‘Divergences’ are cases in which two or more arrows in figures 3 diverge from one melody; according to one criterium only those are counted that arrive at a melody; according to the other criterium any diverging connections are scored (table 1)^9^. Note that some of the parameters in yng are low because his total repertoire is small. In summary, I tentatively interpret the males in the table from left to right as growing in age or at least singing experience.

The connectivity between strophe types may enhance repertoire stability, but only to a limited extent. The receiving strophe is bound to end soon after the transition. It is different with the sequence of strophes in a session: any following strophe is subsequently followed by yet another one, *etcetera*, until the end of the session. This leads us to the next level: how do strophes follow each other? (section c)

#### predictability

However, before entering into this (section c), there is another aspect of strophe structure to be considered, namely the degree of predictability of the course of a strophe once it has begun. To explore this, I analysed, for the four mature birds, those strophe types which occurred at least 9 times in the samples (four types in ♂Sleg, five in ♂tuin, seven in ♂Sand, four in ♂ 1974). Cases where the type melody was actually a second melody were eliminated from the analysis. Table 2 sorts out relatively predictable sequences for variations in the first melody (vars) and for divergences immediately following that melody (divs). (Relatively) predictable are those in which only one possible sequel is realized or where the differences in frequency of diverse sequels is statistically sigificant *(α ≤* 0.05). There is some suggestion in the table of predictability becoming less in the usual order of ♂♂ Sleg, tuin, Sand and 1974, but numbers are small. For all four males taken together about half (45%) of strophe types are completely predictable or almost so by the end of the second element. It must be taken into account, however, that this percentage may further decrease in longer sequences of strophes than the ones available. It is, in fact, this mixture of predictability and variation that creates the network structure of the repertoire. This should be distinguished from the structure of sequences of strophes which we shall explore in the next section.

**Table 2.**
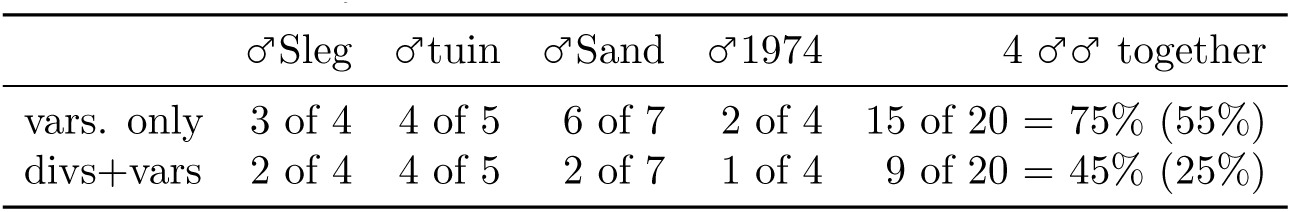
Predictability of the course of a strophe once begun: proportion of relatively predictable sequels (i.c. when differences in frequencies of diverse sequels significant). Four mature males separately and taken together; strophe types occurring ≥ 9× only. Upper row: variations in opening melody only; lower row: combined with divergences in next element (irrespective whether shrill or melody). All cases in which the same melody was not the opening one were eliminated. Last column in parentheses the percentages (of *N* = 20) of connections ending at potential opening melodies only.

### c. Strophe sequences

Some Turdids utter their songs always or almost always in the same order. These are for instance the Mistle Trush (Isaac & Marler, 1963) and Swainson’s Trush (Dobson & Lemon, 1977; Nelson, 1973). Nelson (1989) claims that the succession of song types in the Song Trush is completely random. In search of any regularities in the blackbirds, I plotted the strophes in the order in which they first occur (at the beginning of a record), and found suggestions of repetitions of some strings of types in at least three of the four adult males. By way of example see ♂ 1974 in figures 4A. If for a moment you imagine the figure to represent rain, you may detect a weak but steady wind blowing from left to right. It is as if the singer is turning through a cycle without touching all the same types at every turn. If it is like a 19th century mechanical music box, its pins that trigger the notes are fuzzy. In another metaphor: as if somebody flips through the pages of a song-book with a thumb, stopping here and there but not necessarily at the same pages at every run. Figures 4B-D and 5B-D represent the same procedure for the other three mature males. If the phenomenon is real, one should expect to find more (small) steps down than (larger) steps up in figures 4. Figures 5, which explicate the steps, should then be expected to show a downward trend. This can be tested against a random walk. Table 3 gives values of x^2^ and *p* for all four males. Let me explain the testing procedures and the results in some detail.

**Table 3.** Probabilities of curve of fig. 5 sloping down. Duo’s = pairs, Trio’s = three consecutive strophes; all = all strophes; top = top half of fig. 4 only; −2 = two most frequent strophe types omitted; grf = according to constructed graph (fig. 6). / = up-up; \ = down-down. 1 = series 1; 2 = series 2; 12 = series 1+2. Red: significant at 0.05 level.

**3A. $\sigma$ 1974.**
| | | Duo's ($df = 1$) | | Trio's ($df = 3$) | | / vs \ ($df = 1$) | |
| --- | --- | --- | --- | --- | --- | --- | --- |
| all | 1 | $\chi^2 = 1.5$ | $0.20 < p < 0.30$ | $\chi^2 = 13.168$ | $0.001 < p < 0.01$ | $\chi^2 = 4.5$ | $0.02 < p < 0.05$ |
| | 2 | $\chi^2 = 3.429$ | $0.05 < p < 0.10$ | $\chi^2 = 7.098$ | $0.05 < p < 0.10$ | $\chi^2 = 7.118$ | $0.001 < p < 0.01$ |
| | 12 | $\chi^2 = 4.174$ | $0.02 < p < 0.05$ | $\chi^2 = 18.412$ | $p < 0.001$ | $\chi^2 = 10.796$ | $0.001 < p < 0.01$ |
| top | 1 | $\chi^2 = 1.246$ | $0.20 < p < 0.30$ | $\chi^2 = 3.125$ | $0.30 < p < 0.50$ | $\chi^2 = 2.793$ | $0.05 < p < 0.10$ |
| | 2 | $\chi^2 = 3.125$ | $0.05 < p < 0.10$ | $\chi^2 = 6.032$ | $0.10 < p < 0.20$ | $\chi^2 = 6.231$ | $0.01 < p < 0.02$ |
| | 12 | $\chi^2 = 3.722$ | $0.05 < p < 0.10$ | $\chi^2 = 8.116$ | $0.02 < p < 0.05$ | $\chi^2 = 7.741$ | $0.01 < p < 0.02$ |
| top -2 |  |  |  |  |  |  |  |
| all -2 |  |  |  |  |  |  |  |
| grf | 1 | $\chi^2 = 9.058$ | $0.001 < p < 0.01$ | $\chi^2 = 19.746$ | $p < 0.001$ | $\chi^2 = 14.4$ | $p < 0.001$ |
| | 2 | $\chi^2 = 0.182$ | *n.s.* | $\chi^2 = 1$ | *n.s.* | $\chi^2 = 0.5$ | *n.s.* |
| | 12 | $\chi^2 = 8.011$ | $0.001 < p < 0.01$ | $\chi^2 = 16.977$ | $p < 0.001$ | $\chi^2 = 14.083$ | $p < 0.001$ |

**3B. $\sigma$ Sleg.**
| | | Duo's ($df = 1$) | | Trio's ($df = 3$) | | / vs \ ($df = 1$) | |
| --- | --- | --- | --- | --- | --- | --- | --- |
| all | | $\chi^2 = 1.550$ | $0.20 < p < 0.30$ | $\chi^2 = 9.705$ | $0.02 < p < 0.05$ | $\chi^2 = 4.9$ | $0.02 < p < 0.05$ |
| top | | $\chi^2 = 3.282$ | $0.05 < p < 0.10$ | $\chi^2 = 6.324$ | $0.05 < p < 0.10$ | $\chi^2 = 6.429$ | $0.01 < p < 0.02$ |
| top -2 |  |  |  |  |  |  |  |
| all -2 |  |  |  |  |  |  |  |
| grf | | $\chi^2 = 7.737$ | $0.001 < p < 0.01$ | $\chi^2 = 15.302$ | $0.001 < p < 0.01$ | $\chi^2 = 14.286$ | $p < 0.001$ |

**3C. $\sigma$ tuin**
| | | Duo's ($df = 1$) | | Trio's ($df = 3$) | | / vs \ ($df = 1$) | |
| --- | --- | --- | --- | --- | --- | --- | --- |
| all | | $\chi^2 = 1.052$ | $0.30 < p < 0.50$ | $\chi^2 = 5.819$ | $0.10 < p < 0.20$ | $\chi^2 = 2.814$ | $0.05 < p < 0.10$ |
| top | | $\chi^2 = 0.889$ | $0.30 < p < 0.50$ | $\chi^2 = 5.394$ | $0.10 < p < 0.20$ | $\chi^2 = 2.462$ | $0.05 < p < 0.10$ |
| top -2 | | $\chi^2 = 1.8$ | $0.10 < p < 0.20$ | $\chi^2 = 5.410$ | $0.10 < p < 0.20$ | $\chi^2 = 5.556$ | $0.01 < p < 0.02$ |
| all -2 | | $\chi^2 = 2.997$ | $0.05 < p < 0.10$ | $\chi^2 = 7.158$ | $0.05 < p < 0.10$ | $\chi^2 = 6.125$ | $0.01 < p < 0.02$ |
| grf | | $\chi^2 = 6.722$ | $0.001 < p < 0.01$ | $\chi^2 = 11.985$ | $0.001 < p < 0.01$ | $\chi^2 = 11.765$ | $p < 0.001$ |

**3D. $\sigma$ Sand.**
| | | Duo's ($df = 1$) | | Trio's ($df = 3$) | | / vs \ ($df = 1$) | |
| --- | --- | --- | --- | --- | --- | --- | --- |
| all | 1 | *n.s.* | | $\chi^2 = 7.713$ | $0.05 < p < 0.10$ | $\chi^2 = 1.976$ | $0.10 < p < 0.20$ |
|  | 2 | *n.s.* |  |  | *n.s.* |  | *n.s.* |
| | 12 | *n.s.* | | $\chi^2 = 9.689$ | $0.02 < p < 0.05$ | | *n.s.* |
| top | 1 | *n.s.* | | $\chi^2 = 7.416$ | $0.05 < p < 0.10$ | $\chi^2 = 2.286$ | $0.10 < p < 0.20$ |
| | 2 | *n.s.* | | $\chi^2 = 3.778$ | $0.20 < p < 0.30$ | equal values | |
|  | 12 | *n.s.* |  |  | *n.s.* |  | *n.s.* |
| top -2 | 1 | $\chi^2 = 1.103$ | $0.20 < p < 0.30$ | $\chi^2 = 10.429$ | $0.01 < p < 0.02$ | $\chi^2 = 2.882$ | $0.05 < p < 0.10$ |
| | 2 | $\chi^2 = 0.421$ | *n.s.* | $\chi^2 = 8.333$ | $0.01 < p < 0.02$ | $\chi^2 = 2.778$ | $0.05 < p < 0.10$ |
| | 12 | $\chi^2 = 1.5$ | $0.20 < p < 0.30$ | $\chi^2 = 18.640$ | $0.02 < p < 0.05$ | $\chi^2 = 5.538$ | $0.01 < p < 0.02$ |
| all -2 |  | *n.s.* |  | *n.s.* |  | *n.s.* |  |

**Figure 4.**
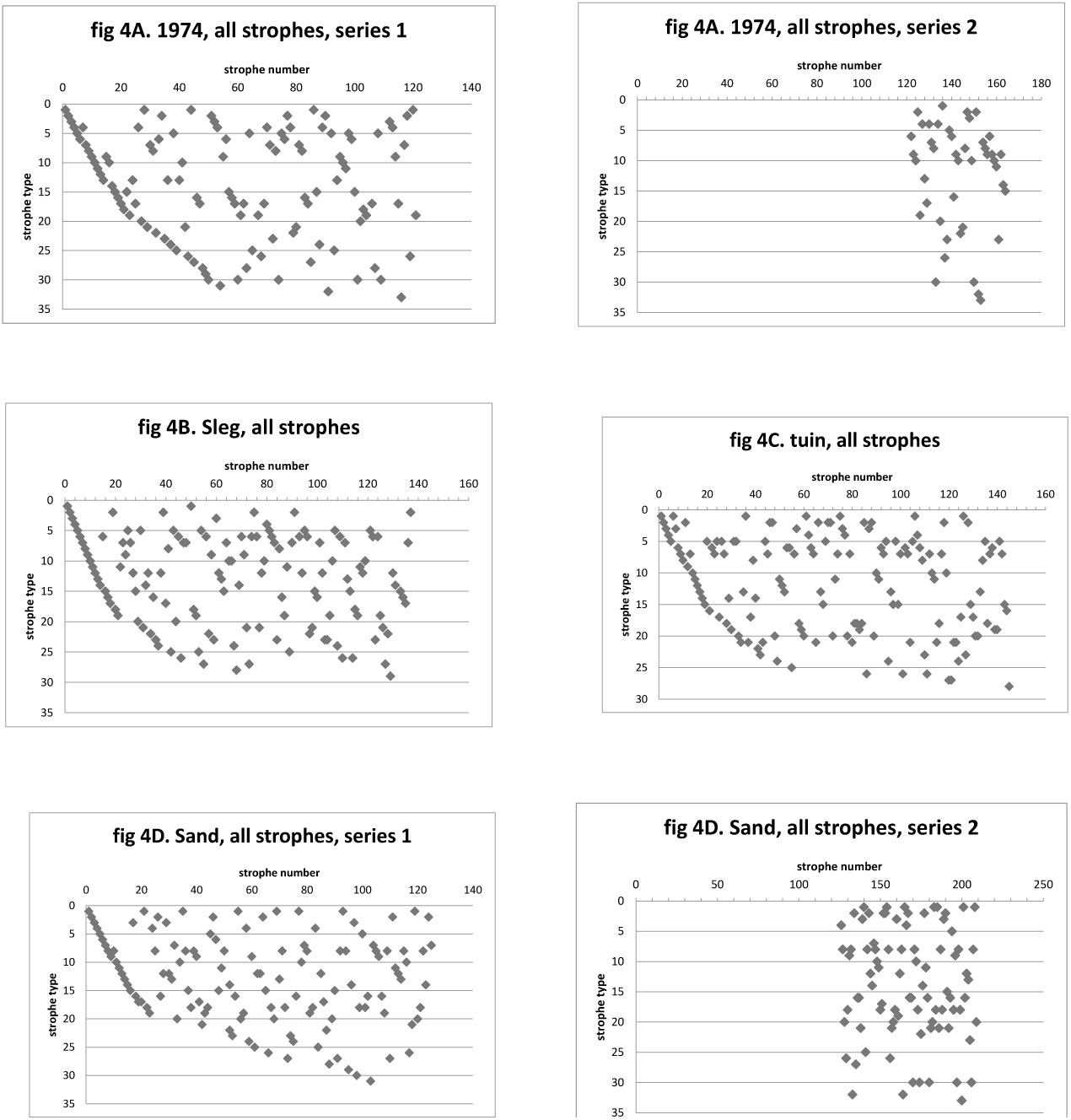
A. All strophes of male 1974 according to their strophe type (ordinate, reading downwards) and their actual order of performance in the records (abscissa). Time (from left to right) measured in strophes, not in seconds. Series 1: before pause in record; series 2: after. B-D. As fig. 4A: males Sleg, tuin and Sand.

**Figure 5.**
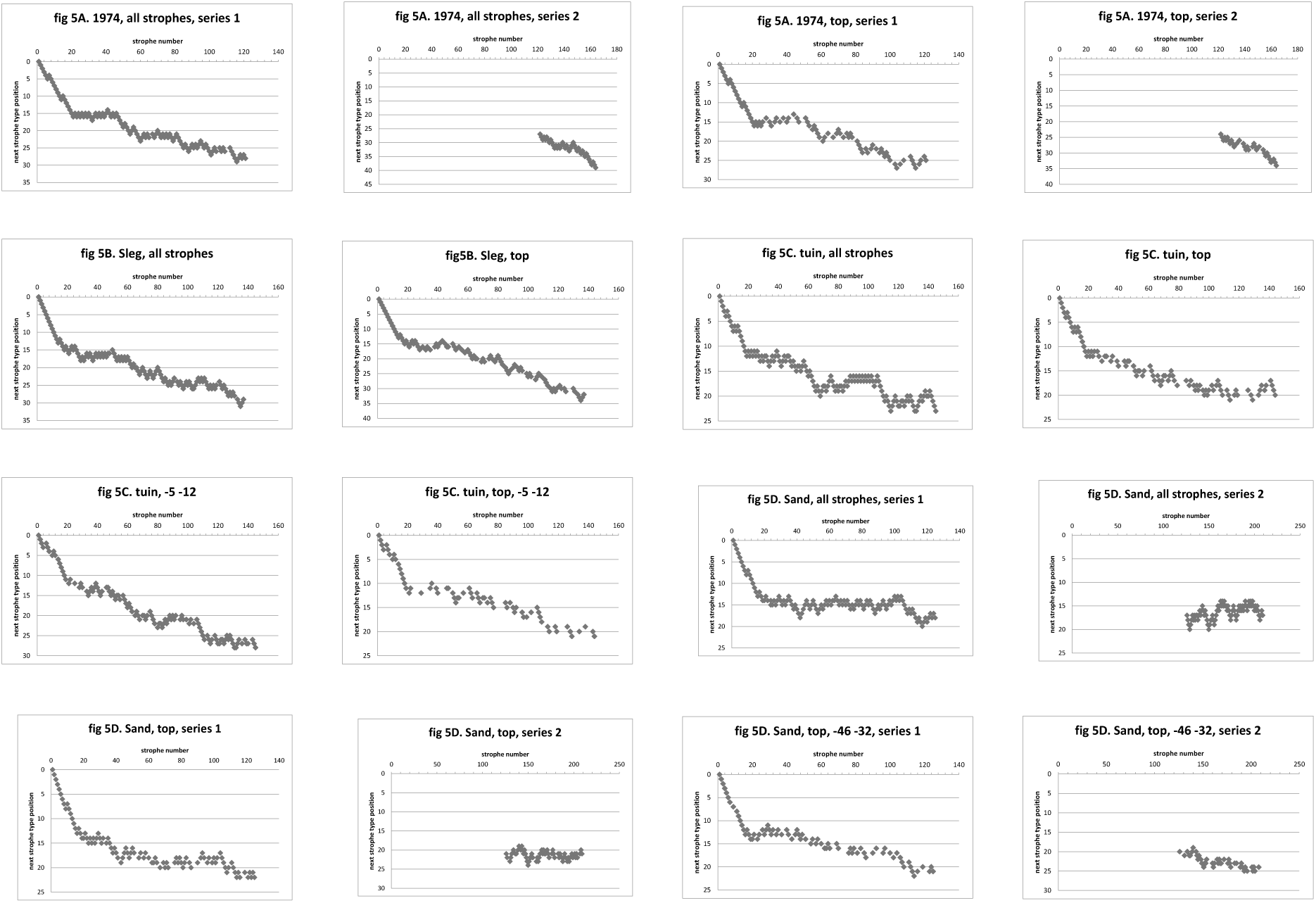
A. Steps up or down (occasionally horizontal = strophe type repetition) of consecutive strophes in fig 4A. All strophes included, respectively (‘top’) only those in top half of fig. 4A (41-23). B-D. As fig. 5A: males Sleg. Tuin and Sand. Respectively: all strophes included, those of top halves of figs. 4B-D only (‘top’), the two most frequent strophe types omitted (−5 −12; resp. −46 −32), or both the latter variations.

The sequences of figure 5A — beginning at strophe number 21 — were broken down into all possible pairs or trio’s. Results of both are presented in Tables 3A-D. I shall only discuss the trio’s (column 2). There are four possibilities: up-up, up-down, down-up and down-down, each of which has a 25% random chance of occurrence. The difference with the real frequencies was χ^2^ tested (column 2 of table 3). In ♂ 1974 the difference is significant for the first series of the graph and for the sum of both series. (Note how close to significance most other results are). The outcomes for the other males are varied. For ♂Sleg the effect is significant, ♂tuin not significant (0.10 *< p <* 0.20) and for ♂ Sand it is almost significant (0.05 *< p <* 0.10) for series 1 and significant for the sum of series 1 and 2.

However, the enemy of the method is that an excess of down-up and up-down trio’s, causing long horizontal parts in the curves, may give rise to false positive outcomes of the tests. So I tested the frequencies of both the former taken together against the other two. The results are significant or almost significant in all four males. This obviously undermines the significances of the former paragraph.

Next, I tried to reduce the number of zig-zagging up and down movements in figures 4. One way to do this is by eliminating the lower half of figure 4, thus suppressing the alternation between new types appearing and earlier ones being repeated. In ♂ 1974 and ♂ Sleg the procedure makes the predominance of up-down and down-up trio’s non-significant. As might be expected, the difference from random is slightly reduced, but still significant in ♂ 1974 and almost significant in ♂Sleg.

As to the other two males, the trick doesn’t work; it even annihilates any significances that were present first. However, in these two males another cause of zig-zagging moves may be found in the comparatively high frequencies of their most frequent strophe types. This was particularly clear in ♂tuin, in which types 5 and 12 happened to be close together on the ordinate (lines 5 and 7 from above, fig. 4C). Therefore, for both ♂tuin and ♂Sand, I eliminated the two most frequent types (46 and 32 in the latter), either together with or without the elimination of the lower half. As a result, the descent of the curves in figure 5 becomes almost significant in ♂tuin (fig 5C) and, in combination with the ‘lower half’ elimination, highly significant in ♂Sand (fig. 5D). Unfortunately, for the latter two males, the excess of up-down and down-up trio’s remains significant despite all eliminations.

Thus, for ♂tuin and ♂ Sand we get stuck in the dilemma whether the significance of the results is due to up-down and down-up sequences being too frequent or the other two sequences *(i.c.* up-up) too rare. The escape from the impasse is in the testing of up-up against down-down trio’s directly (column 3 of tables 3)^10^. These leads to significant results in ♂ 1974 and ♂Sleg, significant results in ♂tuin in at least some of the eliminated forms and even in ♂Sand to a significant result in one of the eliminations.

In addition to the above details must be mentioned that, with few exceptions (♂Sand), all these curves of figures 5 show a downward trend, also if not significant, while none go up.

I think, all this taken together may be interpreted as an asymmetry between the up-up and down-down trends in which a tendency for horizontally suppresses all consistent up-trends (up-up), but not the downward trend. As to what causes the horizontal trend there are at least two possibilities that we have not yet considered. If the model is correct, one has to *expect* one movement up for each cycle. Also: a *randomising* factor would result in a statistically equal number of up and down moves.

We may reconsider the question why the results with ♂ Sand are least clear later *(p.* 18).

#### More complications

Encouraging as the results may be so far, it cannot be the whole story, as the succession of strophes is much more complicated, or more random, than this model. The presence of a random effect is not unlikely (Whitney, 1981b), but it seems worthwhile to see whether the suggested deterministic repetition of strophe types can be made to stand out more clearly. After all, taking the series at the beginning of the record as a standard with which to compare the later ones is somewhat arbitrary. Random effects may have influenced the opening sequence, and strophe types that often follow each other may have been displaced in the same start sequence. Any such hidden relationships are hard to detect with the method applied.

Rather than trying all possible orders, I chose a new approach and plotted all types in a circle and connected with arrows those which immediately follow one another. (see figures 8 on *p.* 15). Then I selected those that occurred at least twice and constructed a simplest graphic representation of these (figs. 6). The results are surprisingly plain: a *single coherent* graph^11^ consisting of a few convergent lines for ♂Sleg, a long series and a slightly more complicated knot or cracknel in ♂tuin and ♂1974, and two star-shaped compartments in ♂Sand, the last one an essentially different configuration from the others. No wonder that ♂Sand had no cycles *(p.* 10)! It may be emphasised that these are transitions, not probabilities of the same. To put one’s faith in transition probabilities is based upon an assumption, namely that frequency of occurrence is a property of the strophe type and not of its position in the network. Evading this assumption and working with the bare transition numbers brought to light an interesting relationship between those strophe types that occupy cross-ways (hubs) in the graphs of fig. 6 and their frequencies as represented in figs. 1. There is even a close correspondence between the frequencies of the disproportionally common types in figs.1 and the degrees of connections (arcs) converging on and diverging from each type (node) in the graphs of figures 6. These correlations are significant (table 4). This finding strongly suggests that the frequencies of strophe types are a function of the number of times they are likely to be met in the pathways of figs. 6.

**Table 4.**
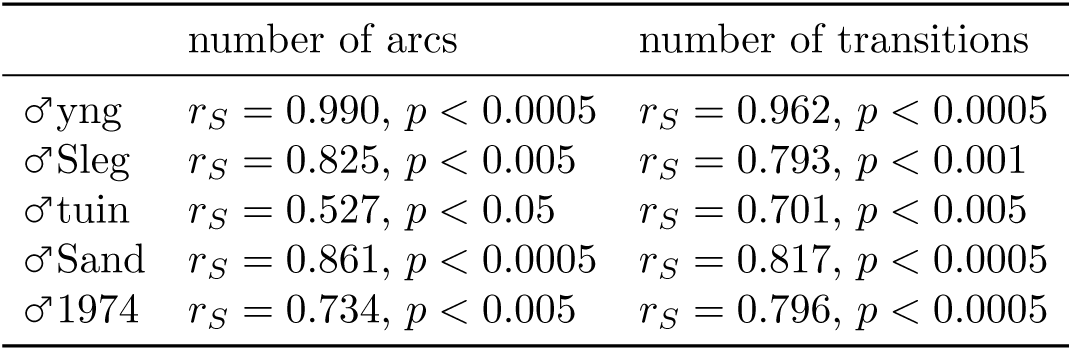
Correlation between degree (= number of arches to and from) of the strophe types in figure 6 and their frequencies in figure 1. Spearman rank correlation onetailed.

**Figure 6.**
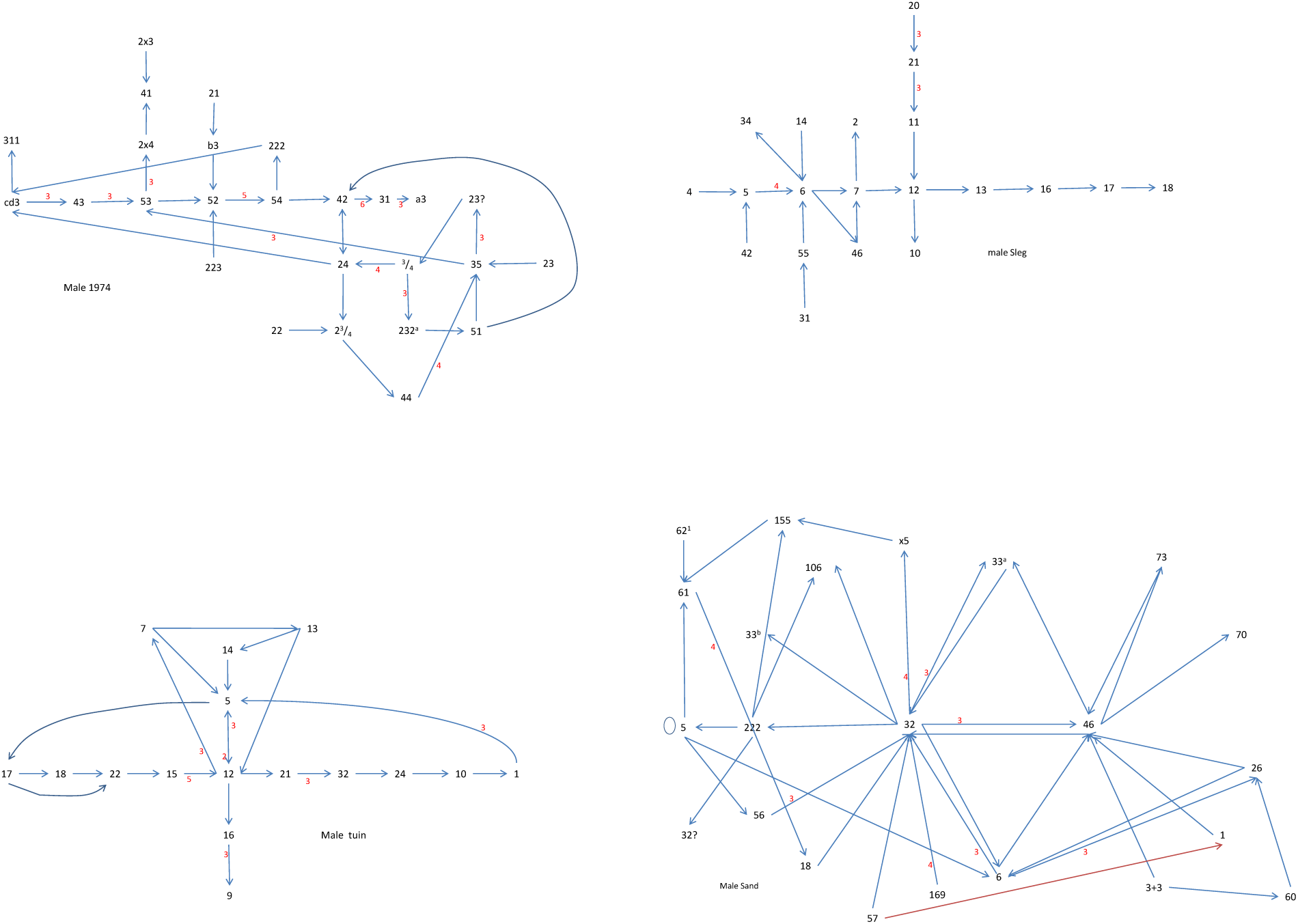
Digraphs of strophe-strophe transitions which occurred at least twice. Males 1974, Sleg, ♂tuin and Sand respectively. Occurrences of more than two indicated with small, red figures.

**Figure 7.**
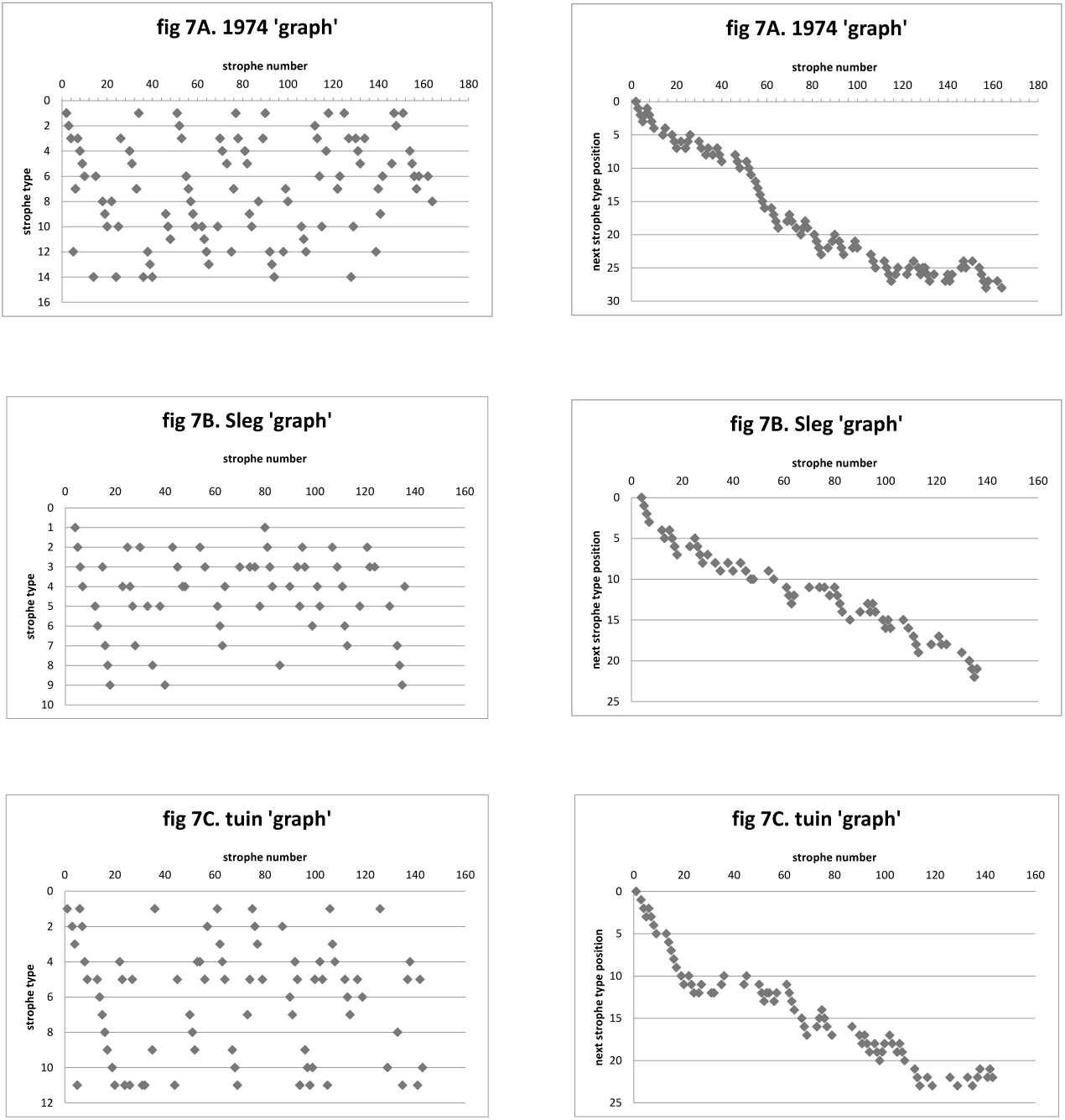
Readings of figures 4A, B and C and figures 5A, B and C with only a selection from the males' repertoires according to the longer chains or cycles of figs. 6. Males 1974, Sleg and tuin. Male Sand (fig. 6D) lacks longer chains and is thus not represented here. ‘Graph’ = derived from the digraphs of figs. 6.

**Figure 8.**
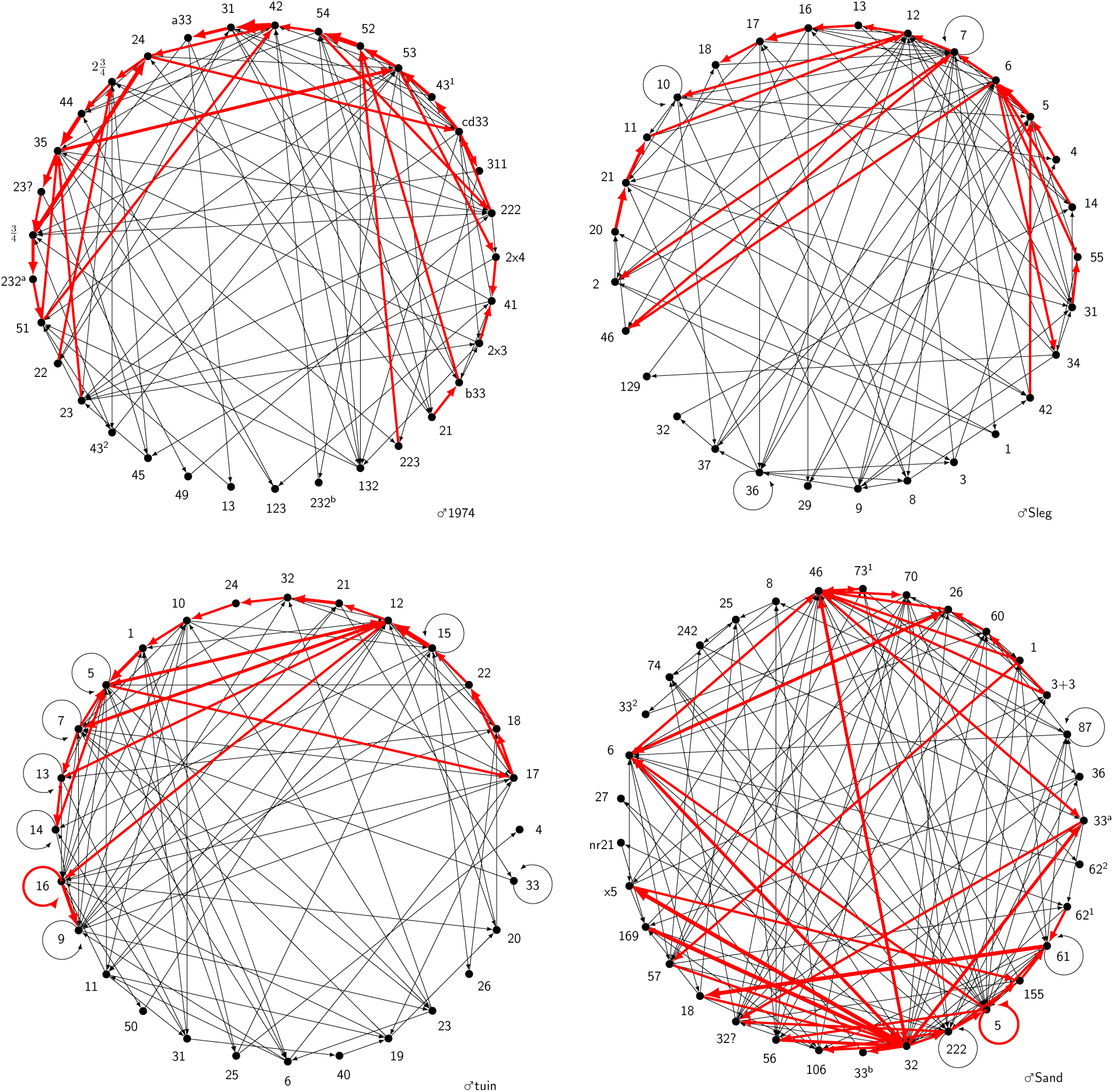
Males 1974, Sleg, ♂tuin and Sand. Digraphs of all strophe-strophe transitions; black = those occurring only once; red = those occurring at least twice — thickness of line corresponding to the frequency of occurrence (max. 6: male 1974, type 42 → 31).

This of course prompts the question where the pathways of graphs 6 come from. There are several possibilities. For instance, the preferred series may hark back to the circumstances in which the strophes were learned, as Todt & Hultsch (1998a) found in the Nightingale. Or there may be relationships between successive strophes as to rhythm, pitch or scale as claimed by Nelson (1973) for Swainson’s Thrush. I have not so far found other key features, but that doesn’t say too much, because the strophes of the blackbird are far more heterogeneous that those of for instance the Varied or Swainson’s Thrush or perhaps even the Nightingale^12^. The way my birds acquired their songs is not known.

#### Unveiling the full cycles

In order to check whether the graphs of figs. 6 represent the cycles of repeated strophe types, I plotted them on the ordinates of graphs like figs. 4. Figs. 7 show the repetitions, standing out more clearly than in the earlier figures. Some come in strings of considerable length, showing that there is more coherence than just some separate pairs of types.

Testing the cyclic coherences in *these* graphs statistically does not prove them to be real, since data were manipulated in order to arrive at them. Nevertheless I added the chi-square tests for comparison in tables 3A-D, bottom row.

#### Order and chaos; the ‘transparency’ of the repertoire

The regularities in strophe type sequences may enhance the conservation of a fair number of characteristic strophe types (those occurring relatively often). Having, however, squeezed the records for deterministic sequences, we may now have a closer look at the more chaotic aspects. Figures 8 visualise the differences between the multiple strophe type transitions and those occurring only once (red and black respectively). In three males the number of single transitions is significantly higher than the total number of transitions of the multiple kind (♂Sleg: 84 *vs* 45; ♂turn: 90 *vs* 54; ♂Sand 118 *vs* 89), so the single ones are worth some attention. Apart from being more common, the single connections are unmistakably more scattered, even in the case of ♂1974, in which the total number of transitions of the multiple kind is higher (*n.s.*): 82 *vs* 74. This scattering may be interpreted as a free accessibility of all strophe types from any other or, in that sense, ‘transparence’of the repertoire. To investigate this further I checked whether the networks of figs. 8 could be qualified as “small-world” networks (p. 4). They do for all four full-grown males. The shortest path length *L* for ♂1974 over 10 random trials was 2.9 (ln33 = 3.50), ♂Sand 2.3 (ln33 = 3.50), ♂♂tuin 2.5 (ln28 = 3.33 and ♂Sleg 2.9 (ln29 = 3.37).

The number of steps needed to reach 90% of the others from a random picked type over 5 trials is shown in Table 5. It may be seen that in all males (hardly more than) 3 steps suffice for 90%, and even as few as two for reaching 50%. Apparently, the connections are of a branching-tree type, which is considered as another indication of “small-world” structure. In other terms: the repertoire is relatively ‘transparent’ as seen from any point in a singing session. This must count as a strong factor in maintaining the full repertoire: no types are likely to be skipped for a long time (and thus perhaps forgotten) in any spontaneous sequence of song.

**Table 5.**
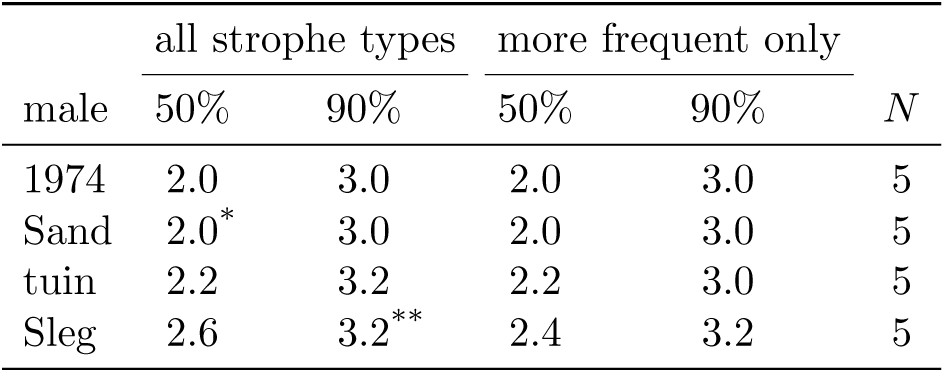
Number of steps, from randomly chosen strophe types, to respectively 50% and 90% of the other types. More frequent: types that occurred more than twice in the samples.

#### Two songbooks?

In males ♂1974 and ♂Sand the graphs (figs 6) consist of two more or less separated pathways: the two ‘stars’ of ♂Sand; the straight sequence and the ‘cracknel’ of ♂1974. The latter seem to cohere mainly via strophe type 42, the former through the direct transitions between 46 and 32 and via types 33a or 6. An interesting question is whether the strophes belonging to either half come in clusters or in random alternation. This was tested with the one-sample runs test (one-tailed; Siegel (1957)) and found to be significantly clustered in most cases. *p* = 0.0455, 0.0024 and 0.0016 for series 1, 2 and 1+2 of ♂Sand; *p* = 0.0021, 0.0868 and < 0.0007 in respectively series 1, 2 and 1+2 of ♂1974). In terms of the above-mentioned metaphor, these birds do not thumb through one song book, but through two in irregular turns! Fascinating thought: could the two syrinx organs account for one book each? Oscine syrinxes are symmetrical and each half can produce sound independent of the other.

## Discussion and Conclusions

The time structures of song in four adult male blackbirds were analysed at three levels: melody variations, divergences within and anastomoses between strophe types, and sequences of the latter. At all of these levels a mixture of preferred paths and divergent scattering was found (table 2 and section c, *p.* 7ff.). The melody variations and the anastomoses between strophe types counteract a breaking up of the repertoire into a collection of rigid types, isolated from each other, and thus work against fragmentation and subsequent loss of repertoire parts. Likewise, the “small-world” nature of the succession ofstrophes keep the whole repertoire ‘transparent’ and accessible from any point in a singing session.

The relationships between successive strophes are probably more important for this effect than the other two, because every strophe *n* +1 which follows n, is itself followed by a strophe *n* + 2 *etc.*, whereas similar chains at the other two levels are soon broken by the limited length of a strophe. Moreover, at least at the middle level the connectivity seems to decrease with individual development, indicating again that it is less important for the conservation of the repertoire. Alternatively, the loss is partly compensated by the succession of a number of strophe types becoming somewhat more stereotyped. From table 5 (♂1974) it appears that this does not need to go at the cost of the general ‘transparency’.

The general ‘transparency’ of the repertoire from any point in the performance is likely to be important too in the facility with which a bird may respond to a neighbour’s song by ‘echoing’ it with its own possibly similar strophe type.

I am not suggesting that all this is the result of evolutionary adaptation. To decide this point more should be known about the development of networks in general and the auditive networks of song birds in particular.

So far we looked at the song structure from the viewpoint of the singer. There is, however, also the aspect of how it is received and responded to by the conspecific listener, whose attention has to be bound by the singer’s voice. It appears to me that also in this respect the mixture of predictable and less predictable sequences may be specially effective. It is this combination of repetition and surprise that is basically effective in human music and we may adopt as a working hypothesis that it is similar in these birds.

We may wonder why the sequence network of ♂Sand is so different from that of the other three adults. One system does not seem to be better than the other where repertoire conservation is concerned. It might be the same with the effects upon the listener, though perhaps the predictability is a little higher, but only as far as the high frequency of occurrence of the two nuclear strophe types is concerned.

One could surmise that the bipolar organisation of ♂Sand’s repertoire (fig. 6D) into two more or less separate ‘communities’ might reduce the ‘transparency’ of his repertoire as a whole, with the particular risk of falling apart in two halves. However, rather surprisingly, ♂Sand’s *L* is lowest of the four males: 2.3 *(p.* 13). I checked whether this low figure could be due to the unintended circumstance that 9 out of 10 trials with ♂Sand were with relatively frequent strophe types *(i.e.* occurring > 2 times). However, a specially made experiment with 10 relatively frequent strophe types of ♂1974 still yielded an *L* of 2.8, almost as high as the 2.9 found earlier and higher than the 2.3 of ♂Sand. Without worrying about significances I think one may safely conclude that ♂Sand’s repertoire is not less ‘transparent’ than those of the others^13^, and thus not more liable to fragmentation and loss.

The state of ♂Sand may be an accident. Among the possible causes of his deformation is that, in his individual experiences, these two strophe types have proved particularly successful with one or more potential mates or rivals. Or perhaps it is the other three males which are atypical. Finally, it may just be that ♂Sand was more than normally disturbed by external stimuli during the recorded session. It is true that many neighbouring males were in full song that evening and during the recordings someone was mowing grass on a hard surface nearby. ♂Sand may have been listening and responding to those other singers, or have been induced by the disturbance to again and again fall back on strophe types learned early in life. Data on more individuals and situations is needed to follow up these fascinating questions.

Finally, it may be mentioned that the networks shown in the figures are only symbolic representations of sequences perceived by the human observer. Their connections, though, must in some way reflect the structure of the singers’ brains, either in the shape of a one-to-one relationship to real neuronal networks or as solutions of activity patterns in circular or more complicated structures as envisaged by Nelson (1973). The fuzziness of the deterministic aspects may be of special interest here.

## Acknowledgements

Many thanks to Mr Yuri Robbers for his encouragement and indispensable help and inspiration at many points throughout the analytic part of the study. I thank Prof. Frank den Hollander for indispensable advice on network theory and calculations, Prof. Dietmar Todt and Dr Henrike Hultsch for reading and commenting on the MS. Mr and Mrs Slegtenhorst and Mrs. Sanders kindly offered the hospitality of their gardens for recording.

## Footnotes

1 *Zoothera naevia*

2 *Turdus migratorius*, *T. philomelos*, *T. merula* and *Luscinia megarhynchos* respectively.

3 *Catharus ustulatus.*

4 Nelson’s data on Swainson’s Thrush can be described as a first order Markov chain. Only, in his case, the relationship between song *n* and song *n* + 1 is always the same as far as it relates to a rough version of a pentatonic scale. This is nothing more mysterious than the inhibition on similar song types in Whitney’s model of the Varied Thrush’s singing, but in an (animal-human) comparative perspective more interesting.

5 *Turdus viscivorus*.

6 Note that the properties described here are valid for very large networks comprising thousands or millions of nodes. They may be assumed to be applicable to small networks such as the present too, but the theory to prove them is a lot more complicated and to date incomplete.

7 In this bird melodies follow one another without intervening shrills. The one exception is marked in the figure.

8 Besides the three second and the seven (potential) first melodies, there are the relatively rare elements 4part and 5part (once each). They consist of one note similar to the (one but) final note of respectively melodies 4 (four notes) and 5 (five notes). Strictly speaking they might be considered as separate types of melodies, bringing the total from 10 to 12. However, I prefer to give the status of ‘melody’ to patterns of more than one note.

9 The numbers of divergences are an approximation. There were too many variations and possible criteria to arrive at an objective counting method.

10 There is some disagreement among statisticians on whether it is justified to test these two fractions against each other. However, the discussion is mainly about cases where significance is very close to *α* (here 0.05), and thus different from most of the results presented in table 3, third column.

11 ‘Digraph’ in the sense of network theory.

12 Successive strophes of a blackbird never begin with the same notes — as has been reported by Todt & Hultsch (1998b) in the Nightingale — except in beginners like ♂yng.

13 Nor is the transparency due to one of the ‘communities’: chosing start and target types from the largest of the two only did not yield any lower *L* than did the whole.

